# Endothelial CXCL12 regulates neovascularization during tissue repair and tumor progression

**DOI:** 10.1101/2020.05.24.113845

**Authors:** Zeshaan N. Maan, Michael S. Hu, Robert Rennert, Janos A. Barrera, Dominik Duscher, Michael Januszyk, Alexander Whittam, Jagannath Padmanabhan, Nicholas Vial, Natalie Ho, Lauren H. Fischer, Johannes Riegler, Joseph C. Wu, Michael T. Longaker, Geoffrey C. Gurtner

**Affiliations:** Department of Surgery, Division of Plastic and Reconstructive Surgery, Stanford University School of Medicine, Stanford, California, USA; Stanford Cardiovascular Institute, Stanford University School of Medicine, Stanford, California, USA

**Author notes:** These authors contributed equally to this work (Michael S. Hu, Robert Rennert). **Corresponding Author:** Geoffrey C. Gurtner, MD, FACS, Professor and Associate Chairman of Surgery, Stanford University School of Medicine, Department of Surgery, Division of Plastic and Reconstructive Surgery, 257 Campus Drive West, Hagey building GK-201, Stanford, CA 94305-5148.

## Abstract

CXC chemokine ligand 12 (CXCL12; stromal cell-derived factor 1 [SDF-1]), primarily known for its role in embryogenesis and hematopoiesis, has also been implicated in tumor biology and neovascularization. However, its specific role and mechanism of action remain poorly understood. We previously demonstrated that CXCL12 expression is Hypoxia-Inducible Factor (HIF)-1 responsive. Here we use a conditional CXCL12 knockout mouse to show that endothelial-specific deletion of CXCL12 (eKO) does not affect embryogenesis, but reduces the survival of ischemic tissue, altering tissue repair and tumor progression. Loss of vascular endothelial CXCL12 disrupts endothelial – fibroblast crosstalk necessary for stromal growth and vascularization. Single-cell gene expression analysis in combination with a parabiosis model reveals a specific population of non-inflammatory circulating cells, defined by genes regulating neovascularization, which is recruited by endothelial CXCL12. These findings indicate an essential role for endothelial CXCL12 expression during the adult neovascular response in tissue injury and tumor progression.

## Introduction

Adult neovascularization (new blood vessel growth in response to ischemia) plays a crucial role in tissue repair and regeneration and influences functional outcomes after injury to the heart, brain, and periphery. The complex biological processes governing neovascularization and tissue repair are exploited by tumor cells (Carmeliet et al., 2011; Hanahan et al., 2011), which typically exist in a setting of relative ischemia (Helmlinger et al., 1997). It is increasingly apparent that tumor survival and progression critically relies on the stromal microenvironment (Li et al., 2003). While numerous studies have explored the importance of a vascularized stroma to both physiologic tissue repair and pathologic tumor development (Valkenburg et al., 2018), an insufficient understanding of the underlying molecular mechanisms has limited the development of effective therapeutics to target this process.

Our laboratory and others have previously described the pivotal role of Chemokine (C-X-C motif) ligand 12 (CXCL12) in the recruitment of circulating cells to hypoxic tissue, secondary to stabilization of Hypoxia-Inducible Factor (HIF)-1 (Ceradini et al., n.d.), and regulation of stem cell microenvironments (Ding et al., 2013; Greenbaum et al., 2013). Interestingly, CXCL12 is also widely expressed in a number of human tumors and has a role in tumor progression and survival (Feig et al., 2013; Orimo et al., 2005; Teicher et al., 2010). These reports suggest a critical role for CXCL12 in both tissue repair and tumor progression, two physiological processes that depend on a delicate orchestration of molecular and cellular factors that govern neovascularization. The mechanisms underlying the influence of CXCL12 on these seemingly disparate processes remains unknown. We hypothesized that interrogating the role of CXCL12 in neovascularization would provide an opportunity to unravel the complex molecular mechanisms that underlie these processes.

Here we use a conditional CXCL12 knockout mouse to identify a critical role for vascular endothelial specific CXCL12 expression in neovascularization and tumor progression. Knockout of endothelial-specific CXCL12 inhibits wound healing and ischemic tissue survival. Surprisingly, loss of endothelial CXCL12 expression completely abrogates tumor growth despite high levels of continued CXCL12 expression by both tumor cells and tumor-associated fibroblasts. We demonstrate that loss of endothelial CXCL12 inhibits fibroblast proliferation, survival, and expression of angiogenic cytokines, clarifying the importance of endothelial-fibroblast cross-talk in this process. Finally, utilizing single-cell gene expression analyses in combination with a parabiosis model, we identify a non-inflammatory progenitor cell subpopulation exclusively recruited to tissue by endothelial CXCL12 signaling. These data reveal a crucial role for vascular endothelial-derived CXCL12 in both tissue repair and tumor progression and uncover a paracrine mechanism governing the formation of vascularized stroma.

## Results

### Vascular endothelial derived CXCL12 does not regulate embryogenesis or vasculogenesis

To specifically assess the role of endothelial CXCL12 during neovascularization, we engineered a floxed allele of *Cxcl12 (Cxcl12 ^loxP/loxP^*) (Figure 1A). *Rosa-creER* and *Tie2-cre* transgenes were utilized to generate tamoxifen-inducible global CXCL12 knockout (gKO) and endothelial-cell-specific (Ding et al., 2013; Greenbaum et al., 2013; Kisanuki et al., 2001) CXCL12 knockout (eKO) mice (Figure 1B), respectively. CXCL12 knockout progeny (*CXCL12^loxP/loxP^; ROSA-creER^+/−^*, *Tie2-cre* ^+/−^) were viable, fertile, produced at expected Mendelian ratios and showed no overt pathologic phenotype. We confirmed DNA recombination upon tamoxifen administration with a subsequent >80% decrease in mRNA and protein expression of CXCL12 (Figure 1C-E). Endothelial specific knockout of CXCL12 was additionally validated used immunostaining (Figure 1F). Prior work has demonstrated that stromal expression of CXCL12 is critical during embryogenesis (Nagasawa et al., 1996), cardiac development (Nagasawa et al., 1996; Tachibana et al., 1998), hematopoiesis (Nagasawa et al., 1996), and organ vascularization (Tachibana et al., 1998). Immunostaining and standard histology were utilized to examine organ development and vascularization in eKO mice and demonstrated normal morphology and vascular pattern (Supplementary Figure 1). This suggests that endothelial CXCL12 does not have a primary role during organogenesis and vascular development.

**Figure 1.**
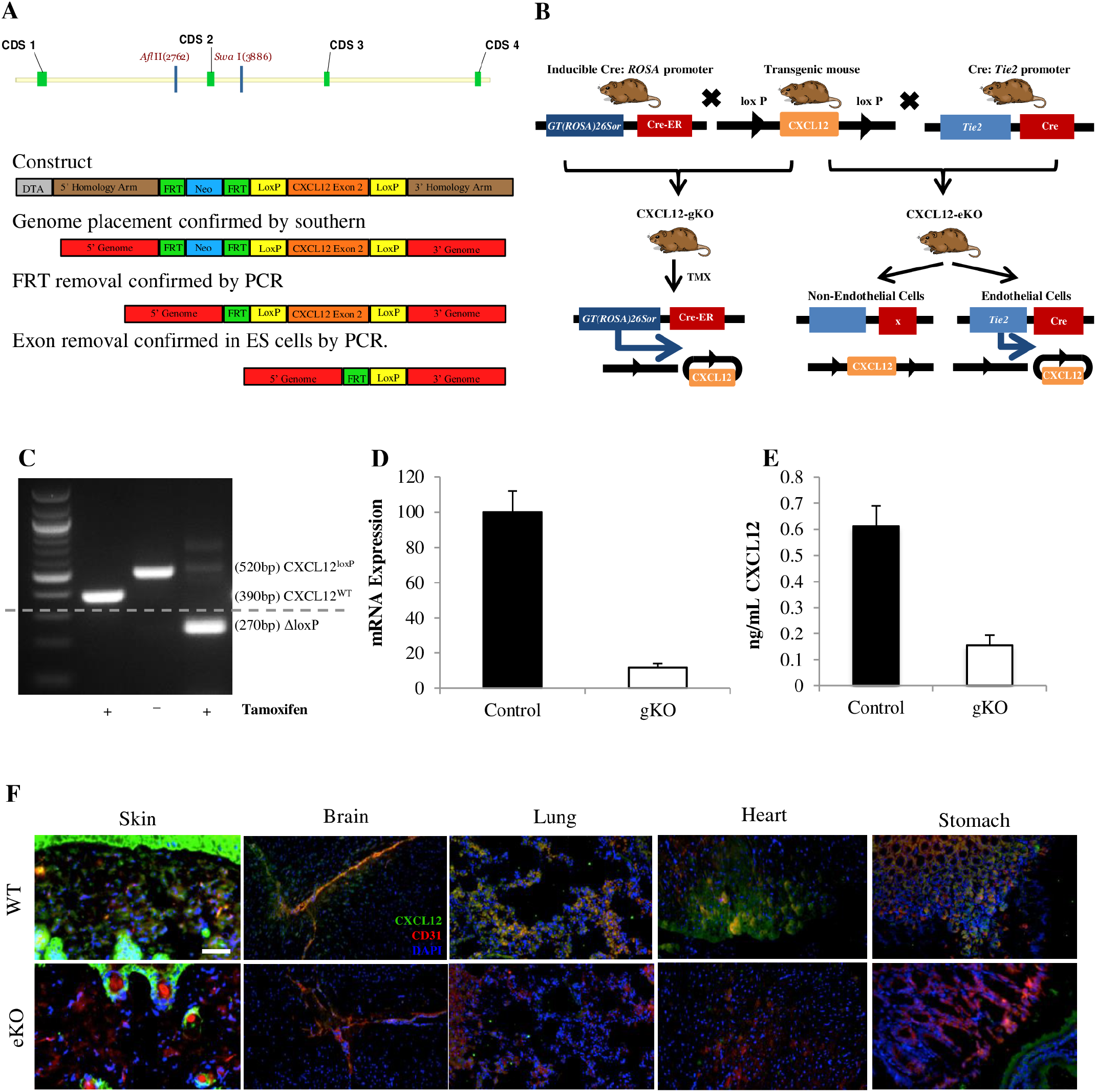
Development and validation of gKO and eKO CXCL12 mice. (A) Schematic of the murine Cxcl12 locus depicting our approach to homologous recombination in embryonic stem cells to generate a floxed Cxcl12 allele. Cxcl12 floxed mice were crossed with mice carrying the *Cre-recombinase* transgene to generate tissue-specific knockout progeny) (B) Schematic of the breeding strategy used to generate tamoxifen inducible global Cxcl12 knockout (gKO) and endothelial-cell specific Cxcl12 knockout (eKO) mice. (C) PCR validation of the presence and recombination of the floxed allele upon exposure of gKO mice to tamoxifen. (D) qRT-PCR of Cxcl12 expression in the skin of tamoxifen induced gKO mice compared to control. (E) ELISA for Cxcl12 levels in the skin of tamoxifen induced gKO mice compared to control. (F) Immunohistochemical analysis showing overlapping expression of CXCL12 and CD31 in wild type but not eKO mice. Images obtained with a Zeiss Axioplan 2 fluorescence microscope, magnification x20, scale bar 200 μm.

### Vascular endothelial CXCL12 critically regulates neovascularization and tissue repair

We then investigated the role of CXCL12 in adult tissue repair using an excisional cutaneous injury model in eKO and gKO mice (Galiano et al., 2004). We found that both eKO and gKO mice showed delayed healing of their wounds compared to floxed control mice (16 days vs. 11 days, respectively) (Figure 2A, B). The impairment in tissue repair in both knockout groups was apparent by the 4th day after injury, as measured by remaining wounded area (Figure 2C). The similar healing times of eKO and gKO mice (Figure 2A-C) suggests that endothelial cells are the critical source of CXCL12 during the repair of injured tissue in adults. Microscopically, the most obvious difference in the healed tissue of eKO mice was a decreased vascular density by immunostaining (Figure 2D, E). We analyzed the injured tissue at earlier time points and found reduced transcription and protein expression of CXCL12 in eKO mice. In addition, VEGF and FGF-2 transcription and protein expression were diminished (Figure 2F-H), suggesting that the CXCL12 signaling modulates other HIF-1 regulated, angiogenic signaling pathways. To confirm the critical role of endothelial CXCL12 signaling during neovascularization, we used a dorsal ischemic skin flap model (Ceradini et al., n.d.), which demonstrated decreased tissue survival in eKO versus wild type mice (Figure 3A-D). Similarly, using a myocardial ischemia model (Patten et al., 1998), in which the left coronary artery is ligated, there was decreased myocardial vessel density in eKO mice (Figure 3E,F).

**Figure 2.**
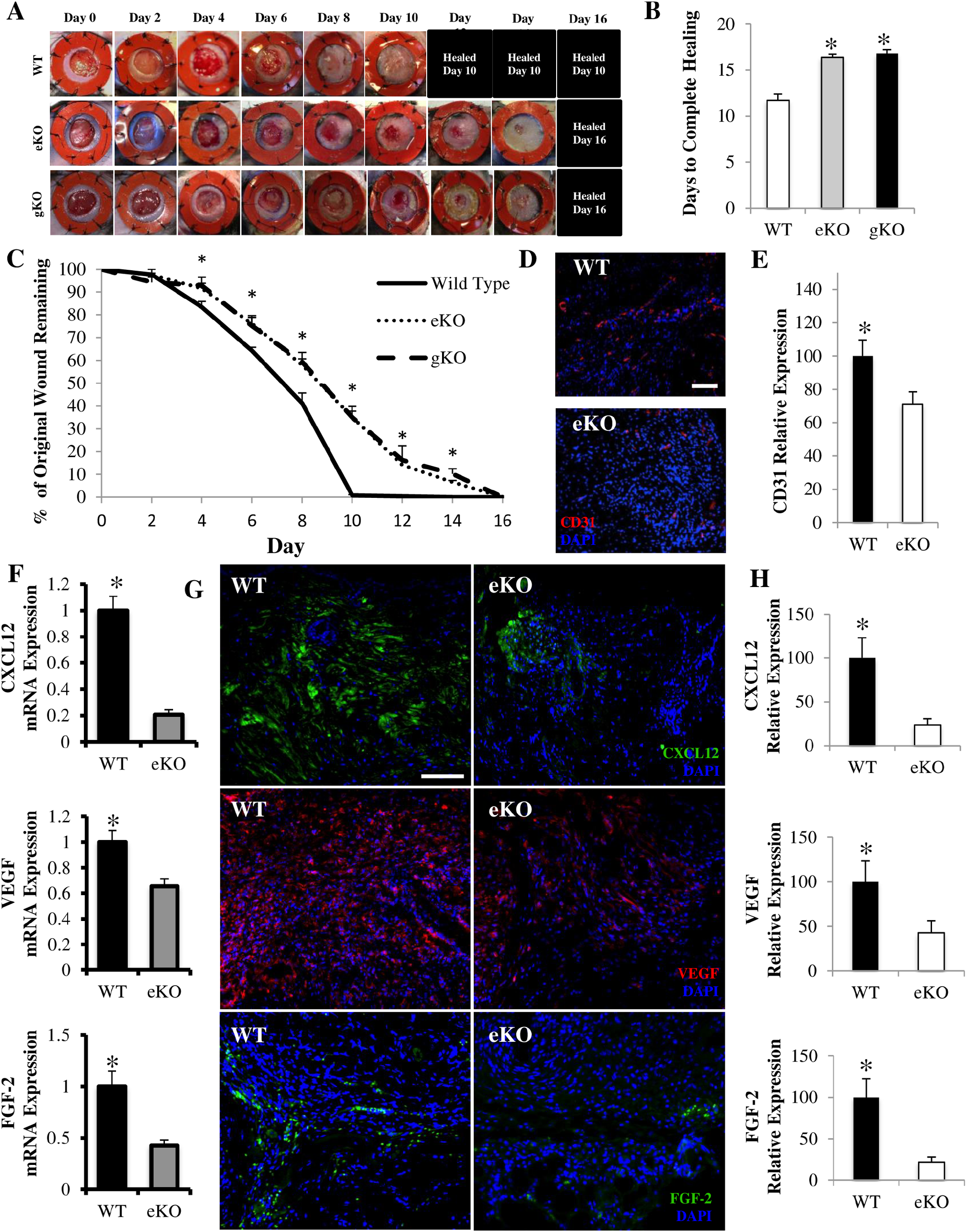
Wound healing relies on endothelial CXCL12. (A) Representative images of similarly delayed healing in endothelial specific (eKO) and global CXCL12 knockout (gKO) mice compared to wild type (WT). (B, C) Both eKO and gKO mice require 16 days to heal compared to 11 days in control. (D, E) Decreased vascular density in fully healed wounds of eKO mice. Images obtained with a Zeiss Axioplan 2 fluorescence microscope, magnification x20, scale bar 200 μm. (F) Decreased expression of Ccxl12 (top), Vegf (middle), and Fgf2 (bottom) in the wounds of eKO mice. (G, H) Selective loss of CXCL12 (top) and decreased levels of VEGF (middle) and FGF-2 (bottom) in eKO mice. Magnification x20, scale bar 200 μm.

**Figure 3.**
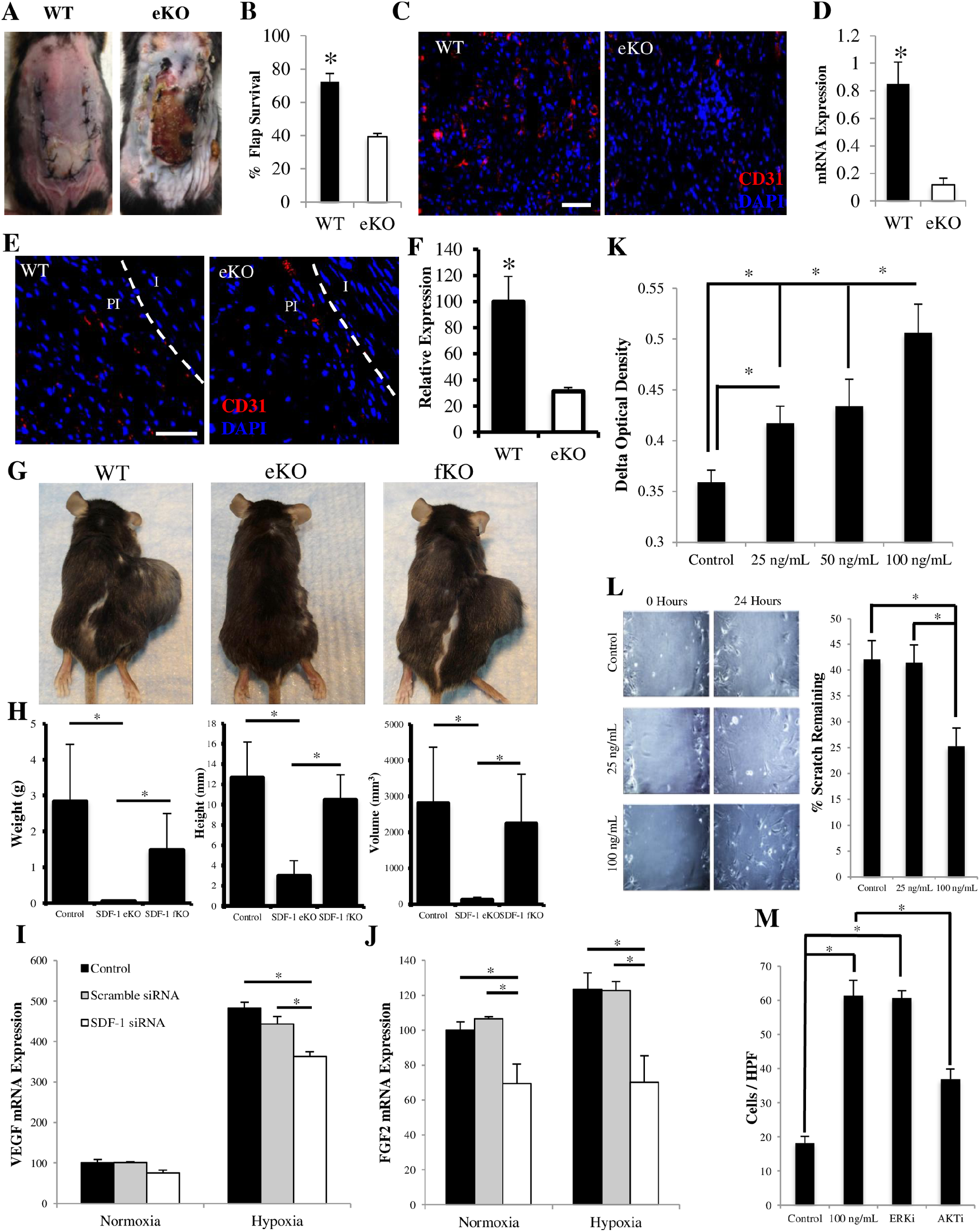
Endothelial CXCL12 regulates neovascularization, stroma formation, and tumor progression. (A, B) Decreased ischemic skin flap survival in eKO mice. (C, D) Decreased vascular density in ischemic tissue of eKO mice. Images obtained with a Zeiss Axioplan 2 fluorescence microscope, magnification x20, scale bar 200 μm. (E, F) Decreased vascular density in peri-infarct zone of eKO myocardium. (PI = peri-infarct zone; I = infarct zone). Magnification x20, scale bar 200 μm. (G, H) Significantly reduced tumor burden in eKO mice. (fKO = fibroblast specific CXCL12 knockout) (I, J) Decreased fibroblast expression of VEGF and FGF-2 in the presence of decreased endothelial expression of CXCL12. (K, L, M) Increased fibroblast proliferation, migratory capacity, and survival (mediated by PI3K/AKT signaling) in response to CXCL12. Images obtained with a Zeiss Axioplan 2 fluorescence microscope, magnification x20, scale bar 200 μm.

### Tumor growth is dependent on host vascular endothelial CXCL12 expression

Cancer cells, which typically exist in the setting of relative ischemia (Helmlinger et al., 1997) appear to rely on the stromal microenvironment for tumor growth, angiogenesis, and invasion (Li et al., 2003). In particular, melanoma has been shown to rely on its surrounding stroma (Flach et al., 2011) and CXCL12, typically derived from cancer cells, has been implicated in tumor progression and survival (Teicher et al., 2010), including in B16 murine melanoma cells (Mendt et al., 2017). To determine if host endothelial CXCL12 had a role in tumor stroma formation and tumor progression, we transplanted B16 murine melanoma cells into control, eKO, and fibroblast-specific (*Col1a2-creER* transgene) (Ubil et al., 2014; Zheng et al., 2002) CXCL12 knockout (fKO) mice. The fKO model was chosen in order to study whether stromal cells themselves produce CXCL12 in sufficient amounts to drive tumor progression, independent of endothelial cell or other circulatory sources. Tumor measurements demonstrated completely abrogated tumor growth only in eKO mice (Figure 3G, H), suggesting a pivotal role for host endothelial CXCL12 in tumor progression.

### Vascular endothelial CXCL12 regulates fibroblast gene expression, proliferation and migration

The establishment of new vascularized tissue, or stroma, requires a coordinated interplay between endothelial cells and fibroblasts (Fukumura et al., 1998; Newman et al., 2011; Stratman et al., 2009). To explore the role of CXCL12 in endothelial – fibroblast cross talk, we used siRNA to target CXCL12 in human microvascular endothelial cells co-cultured with normal human dermal fibroblasts. Decreasing endothelial production of CXCL12 reduced fibroblast expression of VEGF and FGF-2 in response to hypoxia (Figure 3I, J). These data suggest that hypoxia-responsive fibroblast expression of VEGF and FGF-2, which stimulate endothelial cells to proliferate and form new vessels, is in turn reliant on vascular endothelial expression of CXCL12 and provide a possible mechanism for our earlier immunostaining results in injured eKO skin (Figure 2F-H). Additionally, VEGF has been shown to impart drug resistance to tumor endothelial cells within the tumor stroma (Hida et al., 2013), which suggests that endothelial CXCL12 may induce drug resistance in tumors through a paracrine mechanism. Next, we treated fibroblasts with recombinant CXCL12 and demonstrated increased proliferation and migratory capacity (Figure 3K, L). CXCL12 also enhanced the survival of fibroblasts in a low nutrient environment (Figure 3M). It has become increasingly evident that the tumor microenvironment plays a critical role in cancer progression and treatment outcomes. Collectively, these data reveal a novel mechanism by which host endothelial CXCL12 governs the formation of a critical component of this microenvironment and, therefore, tumor progression.

### Vascular endothelial CXCL12 recruits a unique non-inflammatory circulating cell to injured tissue

Because the endothelium is the primary interface between the circulation and the tissue it supplies, and as endothelial cells are crucial for inflammatory cell recruitment (Lawrence et al., 1991; Möhle et al., 1998), we asked whether endothelial specific CXCL12 has a role in the recruitment of non-inflammatory circulating cells that may participate in the neovascularization of hypoxic tissue. Wild type and eKO recipient mice were parabiosed to GFP positive donor mice and cross-circulation was confirmed after 3 weeks using fluorescent imaging and FACS analysis (Figure 4A). Excisional wounds were subsequently created on the dorsum of eKO recipient mice and demonstrated reduced recruitment of mature hematopoietic lineage negative (Ly6C/G, CD45R, TER119, CD4, CD8, and CD11b), GFP positive cells (Figure 4B,C), which would include all types of progenitor cells and exclude inflammatory cells such as neutrophils and lymphocytes. To better characterize the progenitor cells differentially recruited to eKO and control recipient mice, we utilized microfluidic technology to apply a massively parallel single cell transcriptional analysis (SCA) (Supplementary Figure 2, Supplementary Figure 3). Partitional clustering revealed four transcriptionally distinct subpopulations of non-inflammatory progenitor cells in injured tissue. One of these subpopulations (cluster 2) was completely absent in eKO mice, while another was reduced in eKO mice. The remaining two populations were preserved across control and eKO mice (Figure 4D, Supplementary Figure 4, Supplementary Figure 5). The subpopulation completely absent from injured eKO tissue was defined by increased expression of genes associated with progenitor cells, such as Ckit and Mcam, and vascular genes, such as Pecam1, Flt1, Tie1, and Tek. Additionally, this population differentially expressed the cellular adhesion gene Itgb3 (Figure 4E). Pathway analysis software was used to generate a transcriptional network utilizing those genes differentially expressed in this cluster as seed genes. “Inferred” genes included those known to be implicated in cell survival (Erk1/2, Akt, Nfkb complex), neovascularization (Vegf, Pdgfb), and extracellular matrix interactions (Fak). The overall associated biological functions of this network included Cardiovascular System Development and Function, Cellular Movement, and Cell Morphology (Figure 4F). Collectively, these data indicate that endothelial CXCL12 signaling regulates the recruitment of a unique non-inflammatory circulating cell to injured tissue.

**Figure 4.**
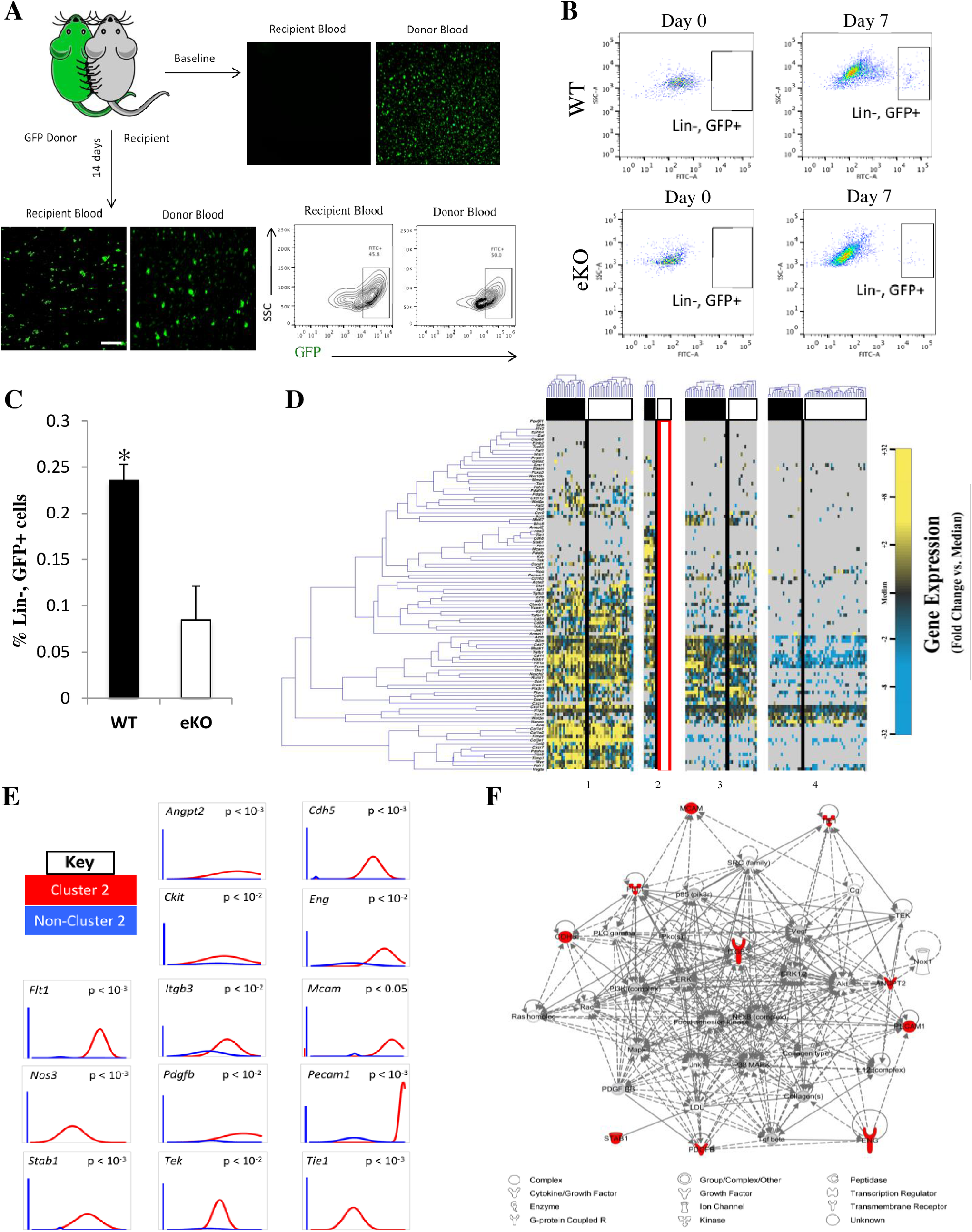
Endothelial CXCL12 regulates recruitment of progenitor cells in ischemic tissue. (A) Parabiosis schema and demonstration of cross-circulation. Images obtained with a Zeiss Axioplan 2 fluorescence microscope, magnification x20, scale bar 200 μm. (B, C) Decreased recruitment of non-inflammatory circulating cells to wounds of eKO mice. (D) Single-cell analysis demonstrating absence of sub-population of non-inflammatory circulating cells from wounds of eKO mice. (E) Differentially expressed genes defining cell population absent from eKO wounds. Left bar for each panel represents fraction of cells that failed to amplify. (F) Top scoring Ingenuity Pathway Analysis (IPA)-constructed transcriptome network based on population defining “seed” genes (E). Seed genes are colored in red, inferred genes in grey.

## Discussion

We describe the differential function of CXCL12 depending on its physiological context and tissue of origin: endothelial expression does not regulate organ development or vascularization during embryogenesis, in contrast to stromal CXCL12 expression (Tachibana et al., 1998), but it does influence the expression of hypoxia-responsive, angiogenic genes and regulates the neovascular response during both the repair of injured tissue and tumor progression. Although the chemotactic effect of CXCL12 in the context of leukocytes has been investigated (Campbell et al., 1998), its physiological function and cellular source in the context of injury had not been previously elucidated. We propose that a distinct population of circulating, non-inflammatory progenitor cells that originate from the bone marrow are exclusively trafficked to injured tissue by CXCL12 mediated angiocrine signaling.

Our findings extend previous work examining the relationship between tumor pathobiology and the mechanisms underlying tissue repair (tumor/wound). Targeting the tumor microenvironment is an emerging paradigm in the management of resistant tumors. Our results demonstrate that host CXCL12 critically regulates the microenvironment independent of tumor derived CXCL12, presenting a potential target for clinical therapy. However, it remains unclear why the selective inactivation of CXCL12 in endothelial cells has such profound effects on tumor growth despite expression of CXCL12 by several other cell types, including tumor endothelial cells. Our findings may also have implications for the development of personalized oncological therapies, as understanding patient-specific biological responses to cancer may be more crucial than tumor profiling. While this study does not exhaustively explore the mechanisms underlying the similar effects of endothelial CXCL12 on tumor growth and tissue repair, there is increasing evidence in the literature of the molecular and cellular similarities between wound healing and tumorigenesis (Arwert et al., 2012; Flier et al., 1986; Schäfer et al., 2008). Further research parsing out potential differences are likely necessary to inform the development of clinical therapies.

## Materials and Methods

### Mice

All transgenic strains had been backcrossed at least ten generations onto a C57BL/6 background (Jackson Laboratories, Bar Harbor, ME). *Rosa–creER, Tie2–cre(14)* and *Col1a2-creER* mice were obtained from the Jackson Laboratory. GFP positive *C57BL/6-Tg(CAG-EGFP)1Osb/J* mice were similarly obtained from Jackson Laboratory. Mice were maintained under standard pathogen-free conditions according to methods approved by the Stanford University Administrative Panel on Laboratory Animal Care (APLAC). All animal experiments were compliant with ethical regulations and approved by the Stanford University APLAC.

### Generation of CXCL12^loxP/loxP^ Mice

A floxed allele of *Cxcl12* was generated by inserting *LoxP* sites flanking exon 2 of *Cxcl12*. FRT sites were inserted flanking the neomycin selection cassette (Figure 1). Generation of targeted embryonic stem cells and blastocyst injections were performed as previously described (Greenbaum et al., 2013). Excision of the neomycin cassette was accomplished through FLP-FRT recombination. Mice were genotyped using PCR primers: *Cxcl12^loxP^* forward, 5’-ACCCATAAATTGAAACATTTGG-3’; *Cxcl12^loxP^* reverse, 5’-TTCTACCACCTGCAGTTTTCC-3’; *Cxcl12^loxP^* recombined, 5’-GGTAAATTTATCGAATTCCGAA-3’.

### Animal Studies

Animal studies were performed with n=5 animals, and repeated at least twice, unless otherwise specified. Mice were 12-16 weeks of age at start of experiments.

#### Murine Excisional Wound Model

Splinted excisional wounds were created as previously described (Galiano et al., 2004). Briefly, using a 6-mm biopsy punch (Integra Miltex, Plainsboro, NJ), two 6-mm full thickness wounds were created on the shaved dorsum of anesthetized mice. In parabiotic pairs, the recipient mouse was only wounded on the non-parabiosed side. A donut-shaped silicone splint (10-mm diameter) was centered on the wound and secured to the skin using an immediate-bonding adhesive (Krazy Glue; Elmer’s Inc., Columbus, Ohio) and 6-0 nylon sutures (Ethicon Inc. Somerville, NJ) to prevent wound contraction. All wounds were covered using an occlusive dressing (Tegaderm; 3M, St. Paul, MN). Following surgery, the mice were placed on warming pads and allowed to fully recover from anesthesia before being returned to the institutional animal facility. Digital photographs were taken at regular intervals and wound area was measured using ImageJ software (NIH) (n = 6 mice). All measurements were performed by a blinded observer. Wound tissue was harvested on post-wounding day 7 and once wounds were completely healed using an 8mm punch biopsy.

#### Murine Ischemic Skin Flap Model

A reproducible model of graded soft tissue ischemia was created on the dorsum of mice as previously described (Valkenburg et al., 2018). Briefly, a full thickness (epidermis, dermis, and underlying adipose tissue) 3-sided peninsular flap (1.25 x 2.5 cm) was created on the shaved dorsum. The flap was elevated from the underlying muscular bed and a 0.13 mm thick silicone sheet was inserted to separate the skin from the underlying tissue. The skin flap was sutured back into place with 6-0 nylon sutures. This model creates a gradient of increasing ischemia from proximal to distal. Digital photographs were taken at regular intervals and necrosed area was measured using ImageJ software (NIH) (n = 6 mice). All measurements were performed by a blinded observer.

#### Murine Myocardial Infarction Model

Ligation of the mid left anterior descending (LAD) artery was performed by a single experienced surgeon (JR) as previously described (Patten et al., 1998). Infarction was confirmed by myocardial blanching and EKG changes. Animals were euthanized and hearts explanted at postoperative day 30 (n=6 mice, repeated once).

#### Murine Parabiosis Model

GFP positive “donor” and either control or eKO “recipient” mice were shaved and anesthetized. Parabiosis surgery was performed as previously described (n=5 mice analyzed) (Bunster et al., 1933; Wagers et al., 2002). Briefly, the corresponding flanks of mice were shaved and disinfected with betadine solution and 70% ethanol. Matching skin incisions were made from the olecranon to the knee joint of each mouse. The skin edges were undermined to create about 1 cm of free skin. 6-0 nylon sutures were used to approximate the dorsal and ventral skin. The skin was over-sewn to protect the suture line. Mice were allowed to recover as described above. Buprenorphine was used for analgesia by subcutaneous injection every 8–12 hours for 48 hours post operation. After three weeks, cross-circulation was confirmed using fluorescent microscopy and FACS analysis (Figure 4A).

#### Murine Tumor Model

A 1 cm incision was created on the right flank of shaved and anesthetized control, eKO, and fKO mice. A 1×1 cm subcutaneous pocket was created and a hydrogel (Wong et al., 2010) seeded with 2.5 x 10^5^ B16-F10 melanoma cells (ATCC CRL-6475) was implanted and the incision closed with 6-0 nylon suture. Measurements for length, width, and height were taken with a digital caliper and tumor volume was calculated using the formula: V = 0.5 x (L x W x H), where V is tumor volume, L is length, W is width, and H is height. Tumor sizes were measured by the same, blinded observer every other day until mice were euthanized at 4 weeks.

### Blood and Skin Analysis

Mononuclear cells from blood were obtained from the buffy coat layer following Ficol-Paque density centrifugation (Jaatinen et al., 2007). Skin samples, including wounded tissue and ischemic skin, were digested and cells isolated as previously described (Suga et al., 2014).

### Histology and Tissue Analysis

For fixation, tissues were placed in 2% paraformaldehyde for 12-16 hours at 4°C. Samples were prepared for embedding by soaking in 30% sucrose in PBS at 4°C for 24 hours. Samples were removed from the sucrose solution and tissue blocks were prepared by embedding in Tissue Tek O.C.T (Sakura Finetek) on dry ice. Frozen blocks were mounted on a MicroM HM550 cryostat (MICROM International GmbH) and 5-8 micron thick sections were transferred to Superfrost/Plus adhesive slides (Fisher & Company, Inc.).

### Immunohistochemistry

For hematoxylin and eosin staining, standardized protocols were used with no modifications. Sections were visualized using Leica DM4000B microscope (Leica Microsystems). Immunostaining on frozen sections was performed using the following primary antibodies: CD31 (Abcam 28364), SDF-1 (Abcam 25117), VEGF (Abcam 52917), and FGF-2 (Abcam 8880). Briefly, slides were fixed in cold acetone (−20°C), and then blocked for 1 hour in 5% goat serum at room temperature followed by incubation with primary antibody for 12-16 hours at 4°C. Slides were then incubated for 1 hour with goat anti-rabbit Alexa Fluor 488 conjugate (Invitrogen A-11034) or 594 conjugate (Invitrogen A-11037). A Zeiss Axioplan 2 fluorescence microscope was used to image the slides (Carl Zeiss, Inc., Thornwood, NY). Quantification of fluorescence was performed by a blinded observer using ImageJ software (NIH).

### Flow Cytometry

All flow cytometry analysis was performed on dissociated wound tissue or blood. Cells were stained by standard protocols with the following fluorescently conjugated antibodies (eBiosciences unless otherwise noted). Lineage analysis was assessed using R-Phycoerythrin (PE)-Cy5-conjugated Ly6C/G (RB6-8C5, Gr-1, myeloid), CD45R (RA3-6B2, B220, B lymphocytes), TER119, CD4, CD8, and CD11b. Cells not stained with these antibodies were incubated with the proper isotype controls or left unstained. Cells were resuspended in FACS buffer and DAPI prior to FACS analysis on a FACSAria II. At least 50,000 events were recorded per sample. Data were analyzed using FlowJo digital fluorescence-activated cell sorting software by a single blinded evaluator (Tree Star Inc, Ashland, OR).

### Quantitative Reverse-Transcription PCR

RNA was isolated using an RNeasy Mini Kit (Qiagen, Hilden, Germany) according to the manufacturer’s instructions. Reverse transcription was performed with 500 ng RNA using the SuperScript III First-Strand Synthesis System (Invitrogen, Carlsbad, CA). qRT-PCR was carried out using TaqMan^®^ Assays-on-Demand™ Gene Expression Products from Applied Biosystems (Foster City, CA, USA): Cxcl12, assay ID Mm00445552_m1; Vegfα, assay ID Mm01281447_m1; Fgf2, assay ID Mm00433287_m1. mRNA expression levels were normalized to B2m expression, assay ID Mm00437762_m1, and presented as relative values.

### *In Vitro* Assays

Human dermal fibroblasts (HdFbs) (Life Technologies C0135C) and human dermal microvascular endothelial cells (HdMVECs) (Life Technologies C01125PA) were purchased and used for *in vitro* assays. All assays were conducted in triplicate unless otherwise stated.

#### Co-Culture

Indirect co-culture experiments were performed using 6-well plates and 0.4 μM pore trans-well inserts. HdFb were seeded in the upper chamber and HdMVECs were seeded in the lower chamber. si-CXCL12 and scrambled-siRNA were purchased form Life Technologies. HMVECs were transfected using Lipofectamine RNAiMAX Reagent (Life Technologies) according to the manufacturer’s protocol before co-culture with HdFb in normoxia and hypoxia as previously described (Ceradini et al., n.d.).

#### Proliferation

Human dermal fibroblasts were plated in 96-well cell-culture plates, 2500 cells/well, in 150 μL of medium with 1% FBS. After 24 hours, fresh media with 1% FBS alone or with varying concentrations of recombinant CXCL12 (25, 50, and 100 ng/mL) were added and after an additional 6 hours BrdU was added and a cell proliferation assay was performed according to the manufacturer’s instructions (Roche Applied Sciences).

#### Migration

Scratch assay was performed as previously described (Wang et al., 2018) on HdFb cultured in 24-well plates with culture medium containing 1% FBS alone or with varying concentrations of recombinant CXCL12 (25 and 100 ng/mL).

#### Survival

HdFb were cultured until 90% confluent in 24-well plates. They were then placed in culture medium containing 1% FBS for 24 hours then cultured in medium containing 0.5% FBS alone, 100 ng/mL recombinant CXCL12, 100 ng/mL recombinant CXCL12 + U0126 (Cell Signaling Technology), or 100 ng/mL recombinant CXCL12 + LY294002 (Cell Signaling Technology). After 72 hours, images of 5 HPFs/well were captured and recorded under phase contrast microscopy and manual cell counts were performed by a blinded observer.

### Microfluidic Single-Cell Gene Expression Analysis

Gene lists were collected from a literature search. Single cell reverse transcription and low cycle pre-amplification were performed as previously described (Glotzbach et al., 2011; Januszyk et al., 2014). Briefly, wound lysate cell suspensions were sorted from *Tie-2^Cre^/CXCL12^loxP/loxP^* and *CXCL12^loxP/loxP^* transgenic mice as single progenitor cells into each well of a 96-well plate using a Becton Dickinson FACSAria flow cytometer (Franklin Lakes, NJ) into 6 μl of lysis buffer and SUPERase-In RNAse inhibitor (Applied Biosystems, Foster City, CA) (N=5 mice). Live/dead gating was performed based on DAPI exclusion. Progenitor cells were defined as previously described. Reverse transcription and low cycle pre-amplification were performed using Cells Direct (Invitrogen) with Taqman assay primer sets (Applied Biosystems) as per the manufacturers specifications. Exon-spanning primers were used where possible to avoid amplification of genomic background. cDNA was loaded onto 96.96 Dynamic Arrays (Fluidigm, South San Francisco, CA) for qPCR amplification using Universal PCR Master Mix (Applied Biosystems) with a uniquely compiled Taqman assay primer set (Supplementary Table 1) as previously described (Glotzbach et al., 2011).

### Statistical Analysis

For comparison between two groups, students t-tests were used with a P-value < 0.05 considered statistically significant. Analysis of single cell data was performed as described previously (Glotzbach et al., 2011; Januszyk et al., 2014; Levi et al., 2011). Briefly, data from all samples were normalized relative to the pooled median expression for each gene and converted to base 2 logarithms. Absolute bounds (+/− 5 cycle thresholds from the median or 32-fold increases/decreases in expression) were set, and non-expressers were assigned to this floor. Clustergrams were then generated using hierarchical clustering (with a ‘complete’ linkage function and Euclidean distance metric) to facilitate data visualization (MATLAB R2011b, MathWorks, Natick, MA).

To detect overlapping patterns within the single cell transcriptional data, k-means clustering was employed using a standard Euclidean distance metric. Accordingly, each cell was assigned membership to a specific cluster as dictated by similarities in expression profiles (minimizing the within-cluster sum of square distances) in MATLAB. Optimally partitioned clusters were then sub-grouped using hierarchical clustering to facilitate visualization of data patterning (Fukumura et al., 1998; Newman et al., 2011). Non-parametric, two-sample Kolmogorov-Smirnov (K-S) tests were used to identify those genes with expression patterns that differed significantly between population clusters and/or groups, following Bonferroni correction for multiple samples using a strict cutoff of p<0.05. For subgroup comparisons, the empirical distribution of cells from each cluster was evaluated against that of the remaining cells in the experiment. Ingenuity Pathway Analysis (IPA, Ingenuity Systems, Redwood City, CA) was used to construct transcriptome networks based on genes that were significantly increased within clusters (including both direct and indirect relationships).

### Data Sharing Statement

Original data may be obtained by e-mail request to the corresponding author. Single cell transcriptional data is available at GEO under accession number GSE146529.

## Acknowledgments

We thank Theresa Carlomagno for administrative support and Yujin Park for help with tissue processing.

