## Supplementary Data for "Endothelial CXCL12 regulates neovascularization during tissue repair and tumor progression"

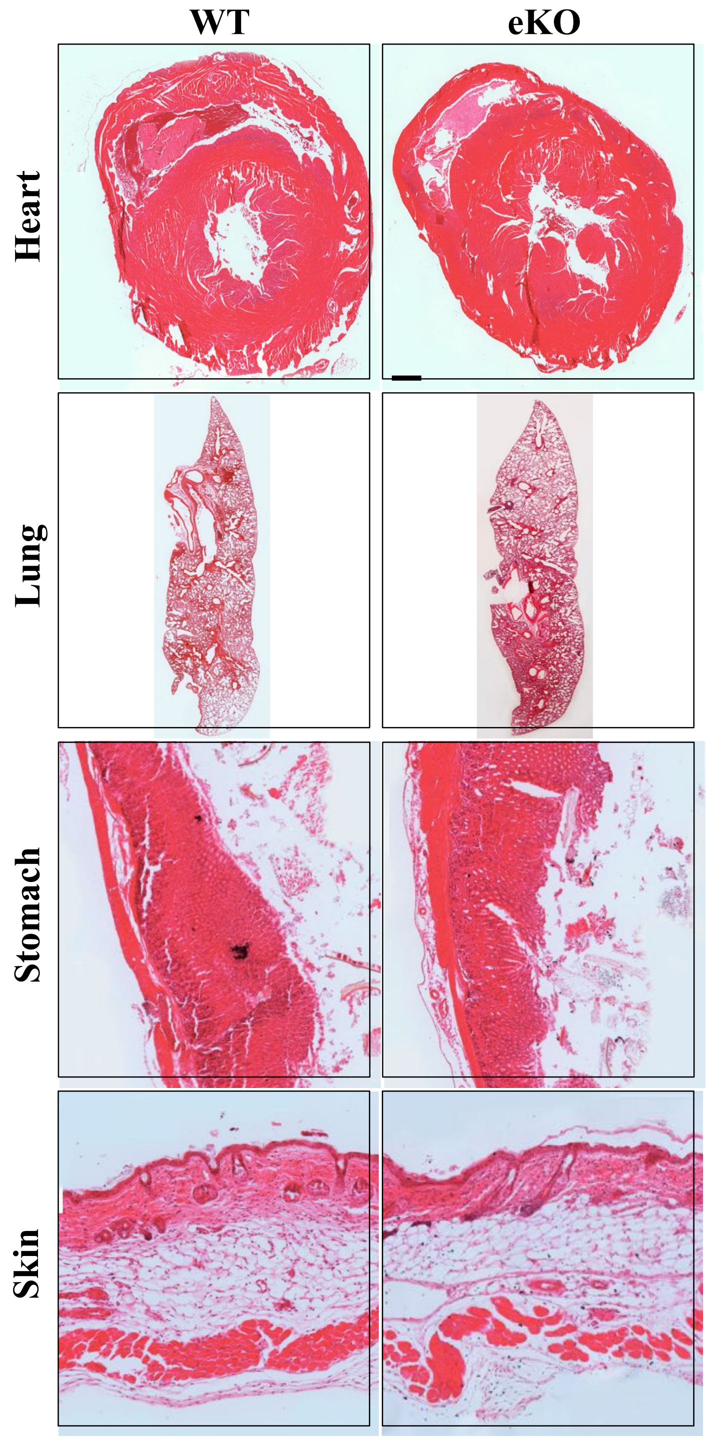


**Supplementary Figure 1. Representative hematoxylin & eosin staining of eKO and wild type organs demonstrating no developmental abnormalities in the absence of endothelial CXCL12**

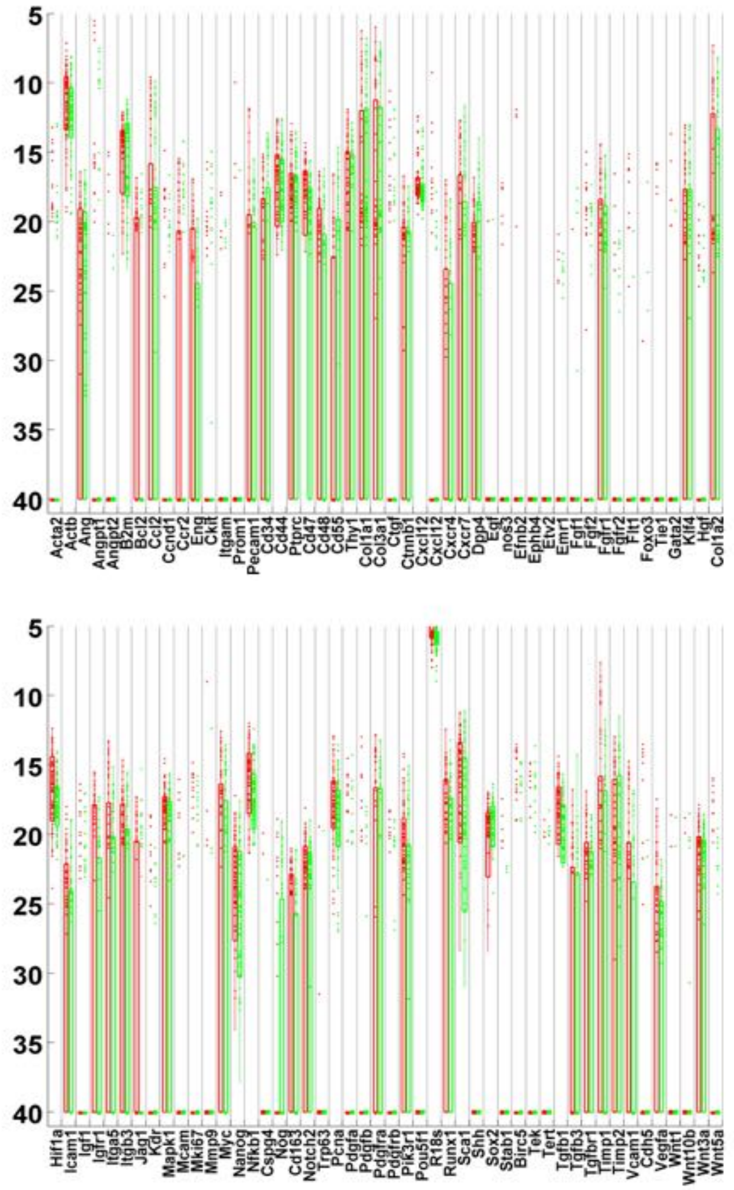


**Supplementary Figure 2. Whisker plots presenting raw qPCR cycle threshold values for each gene across all LIN-, GFP+ cells.** Individual dots represent single gene/cell qPCR reactions, with increased cycle threshold values corresponding to decreased mRNA content. Cycle threshold values of 40 were assigned to all reactions that failed to achieve detectable levels of amplification within 40 qPCR cycles. Cells isolated from wild type and eKO wounds are colored in red and green, respectively. Boxes enclose lower and upper quartiles, and whiskers delimit lowest/highest data points within 1.5 inter-quartile ranges (IQR) or lower/upper quartiles.

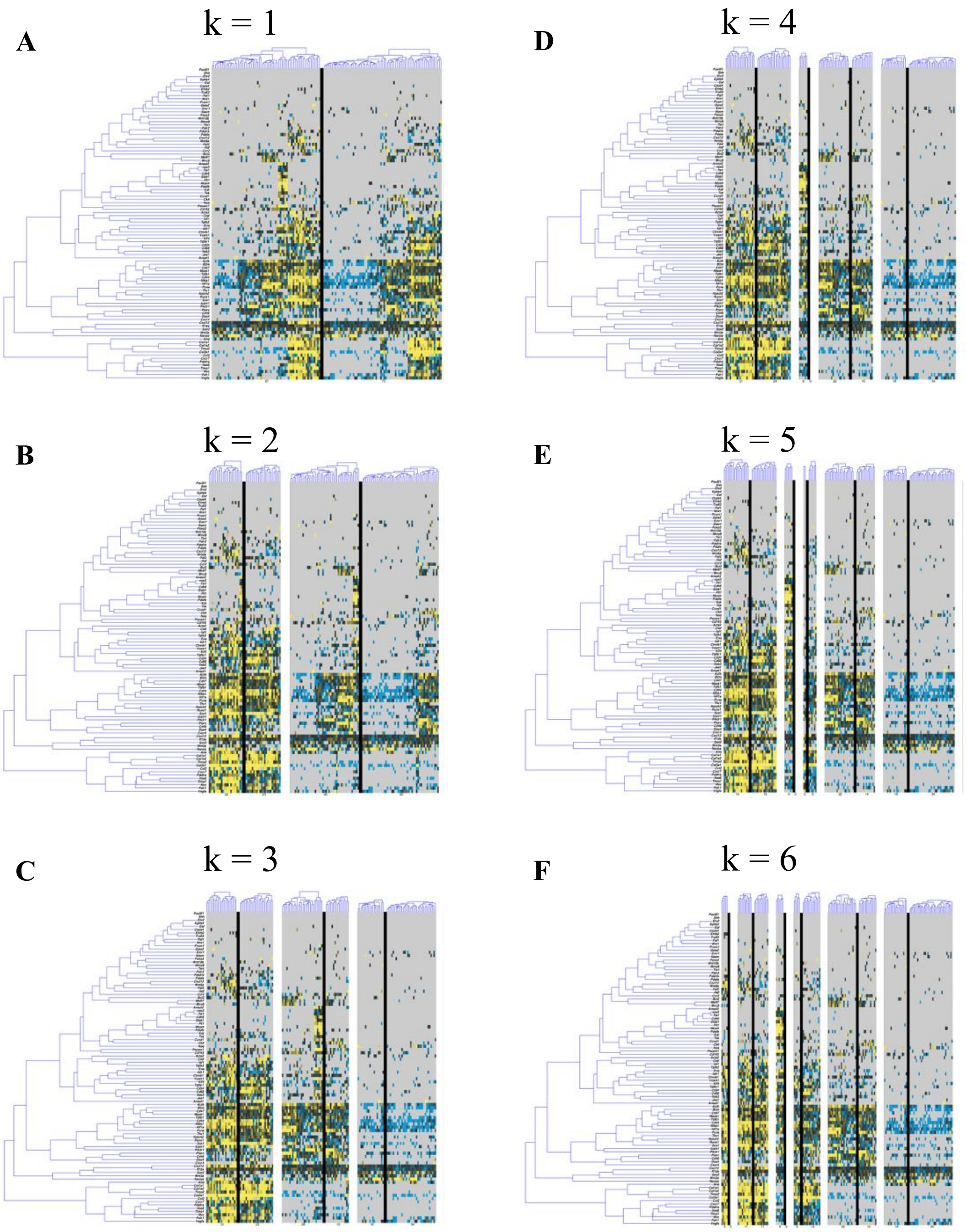


**Supplementary Figure 3. Estimating the number of clusters in single cell gene expression data.** In order to estimate the appropriate number of subgroups present within our dataset, we visualized k=1 to k=6 clusters to estimate the ideal cluster number. The optimal number of clusters is designated as that which maximizes valid clusters without the presence of a "junk" cluster; this corresponded to four clusters in our dataset.

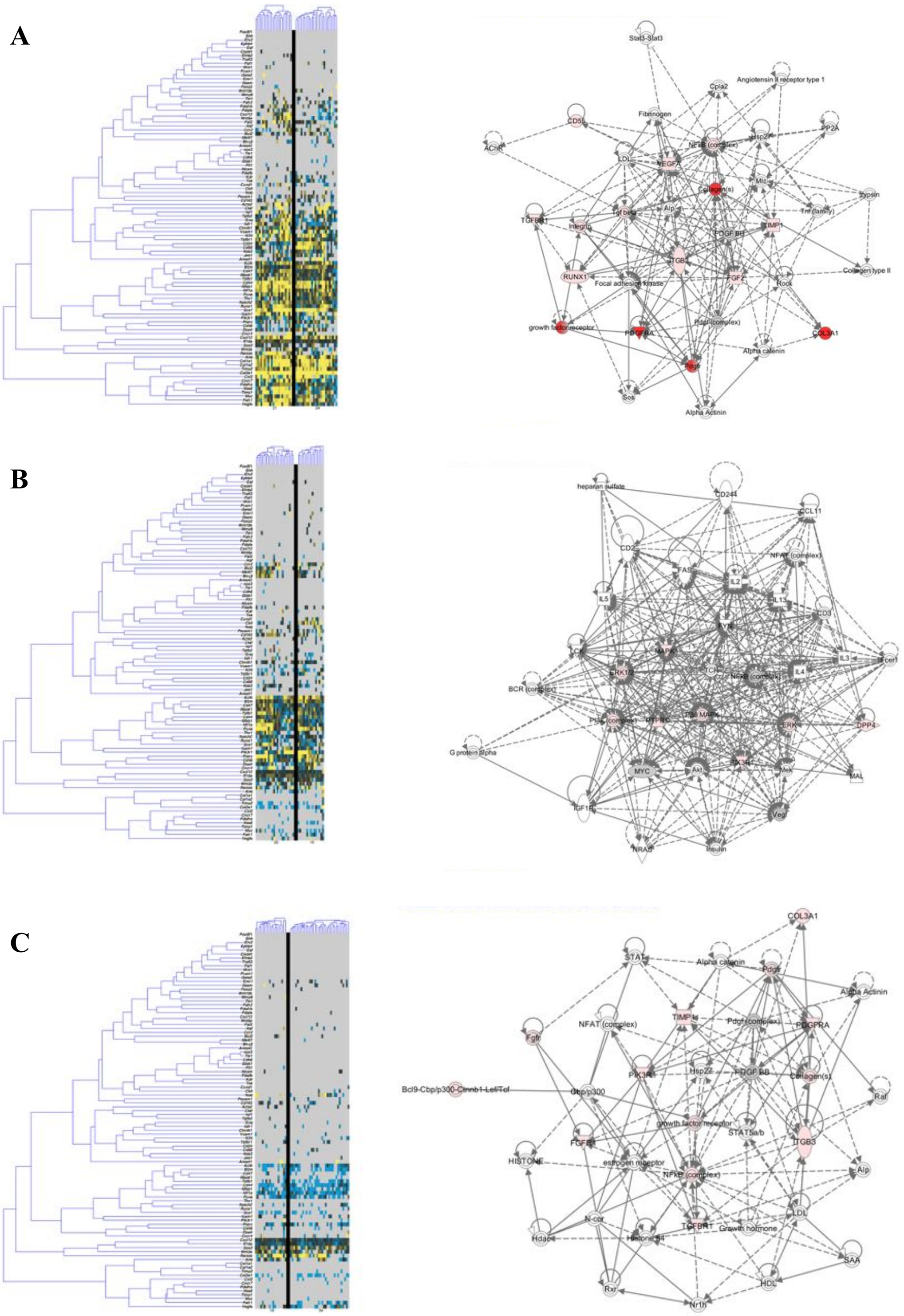


**Supplementary Figure 4. Gene expression in WT (A), gKO (B) and eKO (C) cells as revealed by single cell transcriptome analyses.**

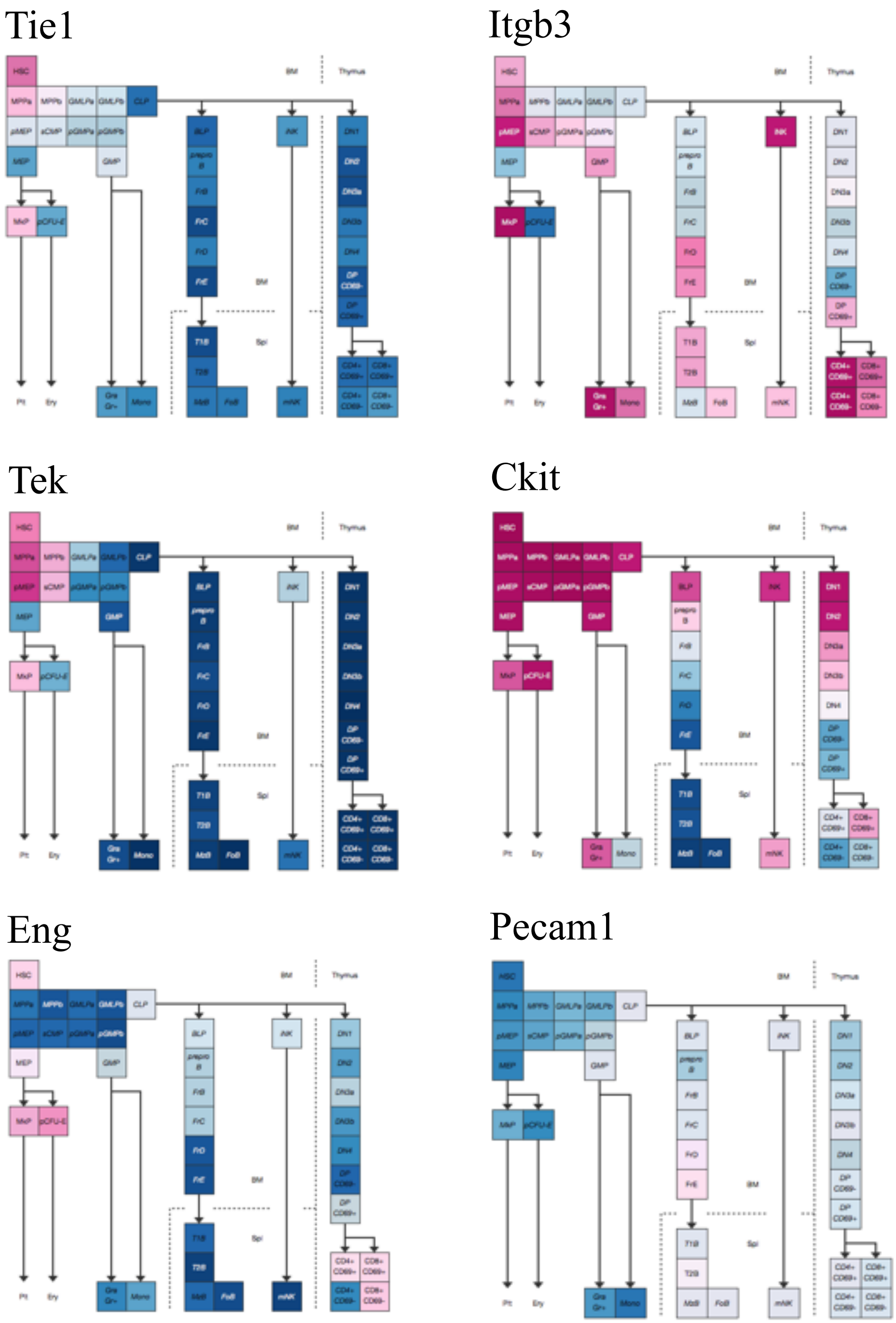


**Supplementary Figure 5. Top scoring Ingenuity Pathway Analysis (IPA)-constructed transcriptome network for the clusters present in both wild type and eKO wounds.** These significant “seed” genes are colored in red to distinguish them from the remaining “inferred” genes in the network.

| Gene Name | Assay ID |
|---|---|
| *Acta2* | Mm01546133_m1 |
| *ActB* | Mm00607939_s1 |
| *Ang* | Mm00833184_s1 |
| *Angpt1* | Mm00456503_m1 |
| *Angpt2* | Mm00545822_m1 |
| *B2m* | Mm00437762_m1 |
| *Bcl2* | Mm00477631_m1 |
| *Birc5* | Mm00599749_m1 |
| *Ccl2* | Mm00441242_m1 |
| *Ccnd1* | Mm00432359_m1 |
| *Ccr2* | Mm01216173_m1 |
| *Cd34* | Mm00519283_m1 |
| *Cd44* | Mm00681165_m1 |
| *Cd47* | Mm00495011_m1 |
| *Cd48* | Mm00455932_m1 |
| *Cd55* | Mm00438377_m1 |
| *Cdh5* | Mm00486938_m1 |
| *Col1a1* | Mm00801666_g1 |
| *Col1a2* | Mm01165187_m1 |
| *Col3a1* | Mm01254473_m1 |
| *Cspg4* | Mm00507257_m1 |
| *Ctgf* | Mm01192932_g1 |
| *Ctnnb1* | Mm00517812_m1 |
| *Cxcl12* | Mm00445553_m1 |
| *Cxcl12* | Mm00457276_m1 |
| *Cxcr4* | Mm01292123_m1 |
| *Cxcr7* | Mm00432610_m1 |
| *Dppiv* | Mm00494538_m1 |
| *Egf* | Mm00438696_m1 |
| *Efnb2* | Mm01215897_m1 |
| *Eng* | Mm00468256_m1 |
| *EphB4* | Mm01201157_m1 |
| *Etv2* | Mm01176580_g1 |
| *Emr1* | Mm00802529_m1 |
| *Fgf1* | Mm00438906_m1 |
| *Fgf2* | Mm00433287_m1 |
| *Fgfr1* | Mm00438930_m1 |
| *Fgfr2* | Mm01269930_m1 |
| *Flt-1* | Mm00438980_m1 |
| *Fox03* | Mm00490673_m1 |
| *Gata2* | Mm00492301_m1 |
| *Hgf* | Mm01135193_m1 |
| *Hif1a* | Mm00468869_m1 |
| *Icam1* | Mm00516023_m1 |
| *Igf1* | Mm00439560_m1 |
| *Igf1r* | Mm00802831_m1 |
| *Itga5* | Mm00439797_m1 |
| *Itgam* | Mm00434464_m1 |

| Gene Name | Assay ID |
|---|---|
| *Itgb3* | Mm00443980_m1 |
| *Jag1* | Mm00496902_m1 |
| *Kdr* | Mm01222421_m1 |
| *Kit* | Mm00445212_m1 |
| *Klf4* | Mm00516104_m1 |
| *Ly6a* | Mm00726565_s1 |
| *Mapk1* | Mm00442479_m1 |
| *Mcam* | Mm00522397_m1 |
| *Mki67* | Mm01278617_m1 |
| *Mmp9* | Mm00442991_m1 |
| *Myc* | Mm00487803_m1 |
| *Nanog* | Mm02019550_s1 |
| *Nfkb1* | Mm00476361_m1 |
| *Nog* | Mm01297833_s1 |
| *Nos3* | Mm00435217_m1 |
| *Notch1* | Mm00435249_m1 |
| *Notch2* | Mm00803077_m1 |
| *Pcna* | Mm00448100_g1 |
| *Pdgfa* | Mm01205760_m1 |
| *Pdgfb* | Mm00440677_m1 |
| *Pdgfra* | Mm00440701_m1 |
| *Pdgfrb* | Mm00435546_m1 |
| *Pecam1* | Mm01242584_m1 |
| *Pik3r1* | Mm00803160_m1 |
| *Pou5f1* | Mm00658129_gH |
| *Prom1* | Mm00477121_m1 |
| *Ptprc* | Mm00448463_m1 |
| *Rn18s* | Mm03928990_g1 |
| *Runx1* | Mm01213405_m1 |
| *Shh* | Mm00436528_m1 |
| *Sox2* | Mm00488369_s1 |
| *Stab1* | Mm00460390_m1 |
| *Tie1* | Mm00441786_m1 |
| *Tek* | Mm01256898_m1 |
| *Tert* | Mm00436931_m1 |
| *Tgfb1* | Mm01178820_m1 |
| *Tgfb3* | Mm00436960_m1 |
| *Tgfbr1* | Mm00436964_m1 |
| *Trp63* | Mm00495788_m1 |
| *Thy1* | Mm00493681_m1 |
| *Timp1* | Mm00441818_m1 |
| *Timp2* | Mm00441825_m1 |
| *Vcam1* | Mm01320970_m1 |
| *Vegfa* | Mm01281447_m1 |
| *Wnt1* | Mm01300555_g1 |
| *Wnt10b* | Mm00442104_m1 |
| *Wnt3a* | Mm01349398_m1 |
| *Wnt5a* | Mm00437347_m1 |

**Supplementary Table 1. Taqman assays used to interrogate gene expression within Lin-/GFP+ cells.** All assays were obtained from Life Technologies (Carlsbad, CA).
